# Reorganization of F-actin nanostructures is required for the late phases of SARS-CoV-2 replication in pulmonary cells

**DOI:** 10.1101/2022.03.08.483451

**Authors:** Jitendriya Swain, Peggy Merida, Karla Rubio, David Bracquemond, Israel Aguilar-Ordoñez, Stefan Günther, Guillermo Barreto, Delphine Muriaux

## Abstract

The severe acute respiratory syndrome coronavirus 2 (SARS-CoV-2) is worldwide the main cause of the COVID-19 pandemic. After infection of human pulmonary cells, intracellular viral replication take place in different cellular compartments resulting in the destruction of the host cells and causing severe respiratory diseases. Although cellular trafficking of SARS-CoV-2 have been explored, little is known about the role of the cytoskeleton during viral replication in pulmonary cells. Here we show that SARS-CoV-2 infection induces dramatic changes of F-actin nanostructures overtime. Ring-like actin nanostructures are surrounding viral intracellular organelles, suggesting a functional interplay between F-actin and viral M clusters during particle assembly. Filopodia-like structures loaded with viruses to neighbour cells suggest these structures as mechanism for cell-to-cell virus transmission. Strikingly, gene expression profile analysis and PKN inhibitor treatments of infected pulmonary cells reveal a major role of alpha-actinins superfamily proteins in SARS-CoV-2 replication. Overall, our results highlight cell actors required for SARS-CoV2 replication that are promises for antiviral targets.

**Teaser:** Impairing regulation of actin filaments inhibits SARS-CoV-2 particle production in human pulmonary cells.

## Introduction

SARS-CoV-2 is worldwide a major public health burden as the main cause of the still ongoing COVID-19 pandemic with almost 244 million confirmed cases in 190 countries and more than 5 million deaths until now (1). SARS-CoV-2 infects mainly human pulmonary cells destroying the target cells and causing severe respiratory diseases with excessive inflammation capable of inducing respiratory failure, multi-organ failure and death (2). SARS-CoV-2 is an enveloped virus with a positive sense, single-stranded RNA genome and belongs to the Beta coronavirus family (3,4,5). After SARS-CoV-2 infection of target cells, intracellular viral replication take place consisting of a series of complex processes (e.g., viral RNA translation, particle packaging, assembly and release) that are tightly orchestrated to one another and often mutually exclusive (reviewed in 6). Viral translation often takes place first in order to create a stock of viral proteins that will serve to assemble the newly made viral particles. The SARS-CoV-2 transcriptome consists of a long unspliced genomic RNA and 9 sub-genomic RNAs that are generated by alternative splicing. After viral RNA translation, once the structural nucleocapsid protein N is produced in the cytosol of infected cells, SARS-CoV-2 assembly continues with the interactions of the N proteins with the unspliced genomic RNA. These interactions lead to a ribonucleoprotein complex that will assemble at the membrane of the Endoplasmic Reticulum – Golgi intermediate Compartment (ERGIC) with the structural proteins transmembrane (M), envelop protein (E) and spike protein (S) (7,8,9). The viral particles, ranging between 90 and 200 nm as recently described (7,8,10,11), will bud from the ERGIC and egress through the secretory pathway. A number of studies over the years have shown that most viruses hijack the cytoskeletal network to fulfill their own replication cycle, which motivated us to perform a detail study on the role of the cytoskeleton during SARS-CoV-2 infection in human host pulmonary cells (12,13,14).

The cytoskeleton is a complex and dynamic network of protein filaments in the cytoplasm of the cells, extending from the cell nucleus to the cell membrane. Its primary function is to give the cell its morphology and mechanical resistance to deformation (15). In addition, the cytoskeleton has been related to many different cellular processes including cell migration, cell signalling, endocytosis, cell division, chromosome segregation, intracellular transport, etc (12, 16,17, 18). It can also build specialized structures, such as flagella, cilia, lamellipodia and podosomes. In eukaryotic cells, the cytoskeleton consists of three main components: microfilaments, intermediate filaments and microtubules, and all these components are rapidly growing or disassembling depending on the requirements of the cell. Microfilaments are mainly composed of linear polymers of G-actin proteins. The G-actin monomer combines to form a polymer which continues to form the actin filament. These actin filament subunits assemble into two chains that intertwine into nanostructures called F-actin chains or fibers (15). F-actin fibers generate force when the growing end of the actin filaments push against a barrier, such as the cell membrane. They also act as tracks for the movement of myosin molecules that affix to the microfilament and “move” along them. Interestingly, the network of F-actin fibers under the plasma membrane can be a carrier for virus entry or transfer from one cell to other (12,17). Although mechanisms of trafficking implicated in SARS-CoV-2 infection have been explored mostly in simian Vero cell lines (19,20), little is known about the participation of F-actin nanostructures during SARS-CoV-2 replication in human pulmonary cells. Here, we implemented confocal and super-resolution 2D and 3D STED microscopy (21) to study the kinetics of M cluster formation during SARS-CoV-2 particle assembly and release, as well as the effects of SARS-CoV-2 infection on the morphology of human pulmonary cells as consequence of intracellular rearrangements of F-actin nanostructures. The human pulmonary A549-hACE2 cells implemented here are a well-established experimental model for SARS-CoV-2 research due to their high susceptibility to SARS-CoV-2 infection, which can be explained by the stably overexpression of the host receptor protein for the viral S protein (human angiotensin-converting enzyme 2, hACE2) and the presence of the the co-receptor (human transmembrane protease serine 2, TMPRSS2) (22). Our results demonstrate that the kinetics of M cluster formation during SARS-CoV-2 particle assembly and release correlate with rearrangements of intracellular F-actin fibers and morphological changes of SARS-CoV-2-infected human pulmonary A549-hACE2 cells. Moreover, we show that the reorganization of F-actin nanostructures is required for SARS-CoV-2 replication in human pulmonary A549-hACE2 cells.

## Results

### M cluster formation during SARS-CoV-2 assembly correlates with changes of F-actin nanostructures and cell morphology

To investigate the kinetics of SARS-CoV-2 assembly, we monitored viral M clusters formation at different time points upon SARS-CoV-2 infection (6 to 77 h post-infection, pi) of human pulmonary A549-hACE2 cells by immunofluorescence confocal microscopy using an anti-COV-2 membrane protein (αM) antibody (Fig. 1A-B, Supplementary Fig. 1A-B and 2). Indeed, human pulmonary A549-hACE2 cells were highly susceptible to SARS-CoV-2 infection (Supplementary Fig. 1 and 2). Further, size quantification of the intracellular M clusters per cell at different time points pi (Fig. 1C, left, and Supplementary Fig. 1B) showed an increase of intracellular M cluster size from a mean of 0.6 μm^2^ (interquartile range, IQR -0.1 -3 μm^2^) at 24h pi to a median of 1.46 μm^2^ (IQR=0.4-15 μm^2^) at 48h pi. Interestingly, the size of intracellular M clusters per cell significantly decreased after 48h pi to 0.77 μm2 (IQR=0.1-9 μm^2^) at 54h pi, 0.72 μm2 (IQR=0.1-8 μm^2^) at 72h pi and 0.64 μm2 (IQR=0.1-7 μm^2^) at 77h pi. Remarkably, the maximal area occupied by intracellular M clusters per cell was 15 μm^2^ at 48h pi, whereas it ranged between 3 and 9 μm2 at the other time points analysed. Quantification of the total intensity of viral M clusters per cell (Figure 1C, right) showed also a significant peak at 48h pi (Mean=8.06 × 10^7^) when compared to the other time points analysed. Supporting these results, RNA-sequencing (RNA-seq) based expression analysis in SARS-CoV-2-infected A549-hACE2 cells showed significant increase of all viral transcripts from 24h pi to 48h pi (Supplementary Fig. 1C), suggesting an increase of unsliced viral genomic RNA during SARS-CoV-2 assembly. Interestingly, we detected intracellular increase of the viral structural proteins N, M and S from 24h pi to 72h pi (Supplementary Fig. 1D), as well as a peak of viral particle release in the cell culture medium at 72h pi (Fig. 1D and Supplementary Fig. 1E). Our results support that during the kinetic of SARS-CoV-2 infection of human pulmonary cells, intracellular SARS-CoV-2 assembly peaks at 48h pi, whereas the release of SARS-CoV-2 particles peaks at 72h pi.

**Fig 1.**
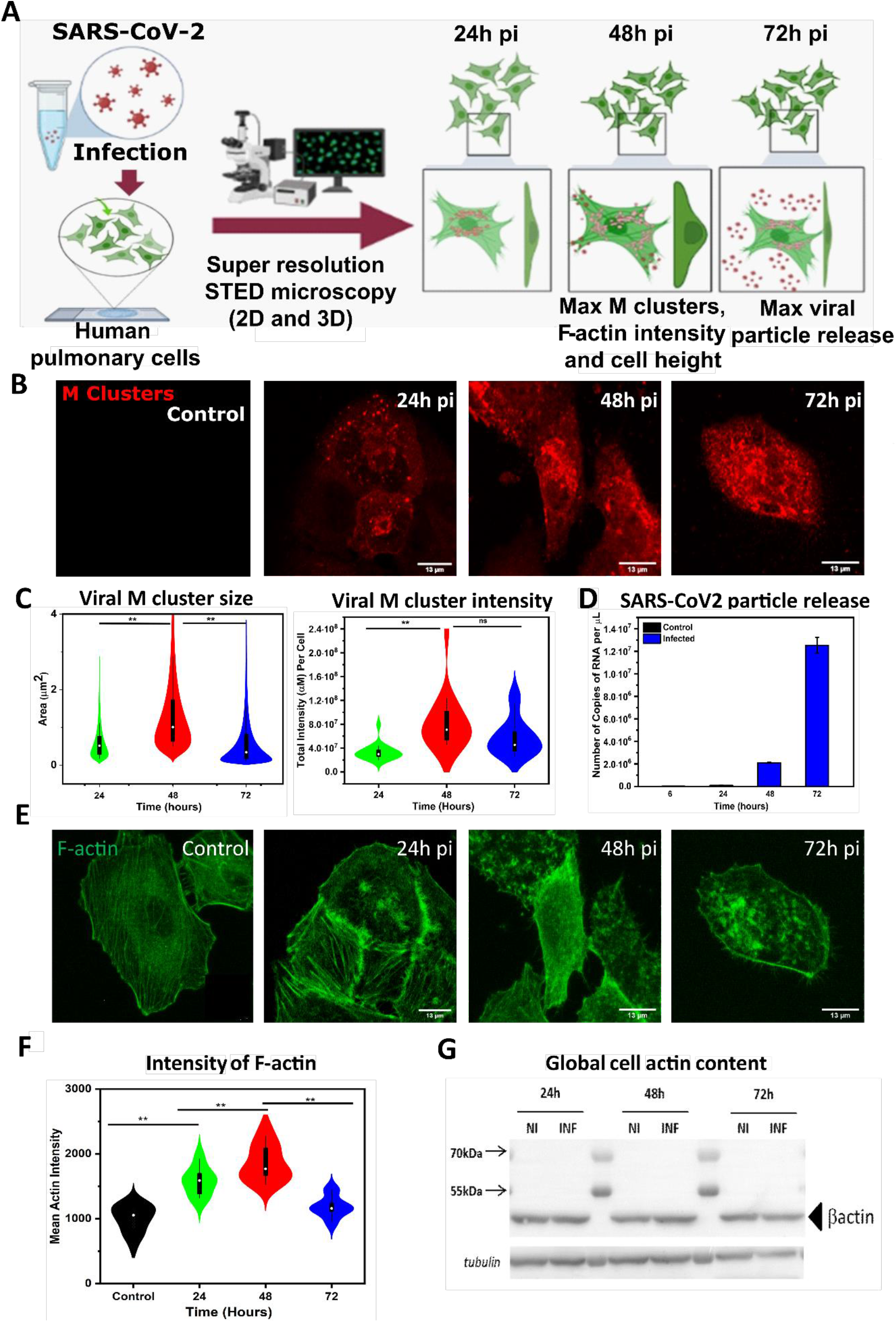
Increase in F-actin content correlated with M clusters of human pulmonary cells upon SARS-CoV-2 infection. Imaging and quantitative analysis of time course changes in viral M clusters area, morphology and mean F-actin intensity of SARS-CoV-2 infected A549-hACE2 cells. A549-hACE2 cells were fixed at 0h, 24h, 48h, and 72h post infection and processed for immunofluorescence. SARS membrane protein anti-M rabbit antibody and a secondary antibody Alexa Fluor 568 (in red) and for F-actin Phalloidin Alexa Fluor 488 were used for confocal microscopy. (A) Schematic representation and confocal images of viral clusters and F-actin with different time post infection 0h to 72h. (B) Images for changes in viral M clusters size at different time post infection. (C) Plot for viral M clusters size and viral M clusters intensity at different time post infection. (D) Plot for Number of copies of RNA/μL in the supernatant of infected cells with different time post infection. (E) Images for changes in F-actin intensity per cell at different time post infection. (F) Plot for mean F-actin intensity with or without infection at different time of post infection. (G) Western blot data for global actin content of non-infected and infected cell at different time point of infection. All F-actin intensity of infected and non-infected cells are calculated from Z-projection images. A number of 20 < *n* <50 cells were analyzed from at least 3 independent experiments. Statistical significant analysis were evaluated using one‐way ANOVA tests. \*\**p* < 0.05.

We also monitored the effect of SARS-CoV-2 infection on the cytoskeleton of A549-hACE2 cells by confocal microscopy at different time points pi after labelling F-actin by phalloidin stain (Figure 1E and Supplementary Fig. 2). We observed changes in the distribution of F-actin in A549-hACE2 cells at different time points SARS-CoV-2 pi. *E*.*g*., F-actin stress fibers were visible in non-infected control cells and disappeared 24h pi, whereas other F-actin structures, such as elongated filopodia, appeared 24h pi. Further, quantification of F-actin intensity implementing Z-stack projection images (Fig. 1F) revealed significant increase of F-actin intensity from 999 a.u. at 0h pi to 1572 a.u. at 24h pi and 1857 a.u. at 48h pi, whereas F-actin intensity decreased to 1184 a.u. at 72h pi. Interestingly, we did not detect significant changes in the total actin content of the cell by western blot analysis (WB) of proteins extracts (Figure 1G). Our results indicate that SARS-CoV-2 infection induced extensive reorganization of intracellular actin fibers without significantly affecting total actin levels.

Further analysis of the morphological changes that we observed in A549-hACE2 cells upon SARS-CoV-2 infection (Fig. 2A-B, Supplementary Movies 1 and 2) revealed significant increase of cell height from 2.96 μm (IQR=2.2 -3.9 μm) at 0h pi to 3.75 μm (IQR=2.2-4.72 μm) at 24h pi and 4.89 μm (IQR= 3.4-6 μm) at 48h pi, whereas cell height decreased to 3.29 μm (IQR=2.4-4.8 μm) at 72h pi. In contrast to cell height, cell surface and cell volume showed the opposite effects upon SARS-CoV-2 infection with decreases values above 24h pi (Fig. 2C-D), suggesting a contraction of the A549-hACE2 cells at this time point. Summarizing, our results indicate that the kinetics of M cluster formation during SARS-CoV-2 particle assembly and release correlate with rearrangements of intracellular F-actin fibers and morphological changes of SARS-CoV-2-infected human pulmonary A549-hACE2 cells.

**Fig 2.**
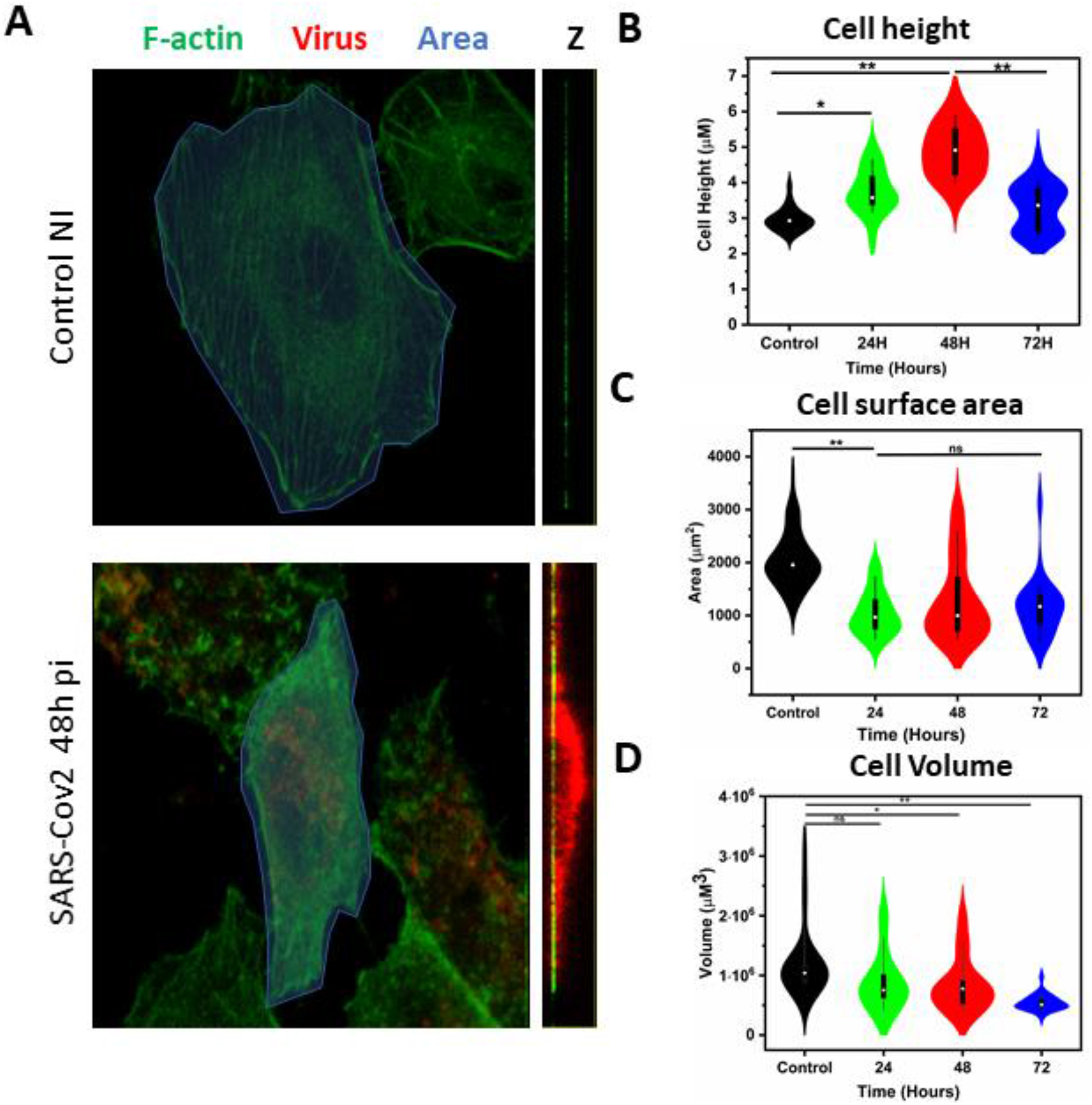
Morphological changes of human pulmonary cells upon SARS-CoV-2 infection. Imaging and quantitative analysis of time course changes in morphology of SARS-CoV-2 infected A549-hACE2 cells. A549-hACE2 cells were fixed at 0h, 24h, 48h, and 72h post infection and processed for immunofluorescence. SARS membrane protein anti-M rabbit antibody and a secondary antibody Alexa Fluor 568 (in red) and for F-actin Phalloidin Alexa Fluor 488 were used for confocal microscopy. (A) Confocal images of Control and Infected A549-hACE2 cells (48h post infection. (B) Plot for cell height. (C) Plot for surface area of the cell. (D) plot for cell volume. All cell surface area, height, volume of infected and non-infected cells are calculated from actin and M protein spreading at XY and Z direction at different time of post infection, A number of 20 < *n* <50 cells were analyzed from at least 3 independent experiments. Statistically significant analysis was evaluated using one‐way ANOVA tests. \*\**p* < 0.05.

### Reorganization of F-actin nanostructures in SARS-CoV-2 infected pulmonary cells

Since we detected a maximum of M cluster formation during SARS-CoV-2 assembly at 48h pi, we focused our further analysis on this specific time point and optimized experimental conditions to achieve a higher resolution for imaging of F-actin structures using super-resolution 2D STED microscopy in non-infected and SARS-CoV-2 infected A549-hACE2 cells (Figure 3A), thereby increasing resolution below 70nm by 10 times as compared to confocal microscopy and being able to observe dual colour structures. To quantify F-actin rearrangements, we have analyzed the orientation angle of actin fibers from STED images (as in 18). We clearly observed the parallel orientation of actin stress fibers in non-infected cells, whereas actin stress fibers were not visible in SARS-CoV-2 infected cells at 48h pi (Fig. 3A). The color map of F-actin orientation and data for distribution of orientation angle, suggested a significant rearrangement of F-actin network with protrusion of filaments at the cell plasma membrane (Fig. 3A-B). The major possible orientation angle of F-actin fibers in non-infected cells was significantly lower (around 90 and -90 degree) than in SARS-CoV-2 infected cells at 48h pi (Figure 2 B). We also detected an increase in the random orientation angle after infection, probably due to F-actin rearrangement, distortion or reorganization. To investigate the potential formation of intracellular self-organizing F-actin structures in SARS-CoV-2 infected A549-hACE2 cells, we monitored the organization of the F-actin cytoskeleton at 48h pi using super-resolution 2D microscopy and detected in the infected cells F-actin structures resembling intracellular “actin rings” (Fig. 3C). The diameter of the observed “actin rings” ranged between 0.5 and 2.5 μm with a mean of 1.03 μm and a standard deviation (STD) of 0.35 μm (Fig. 3D, top). However, detection of viral M clusters around intracellular “actin rings” by 2D STED microscopy was limited by signal saturation in Z direction. To bypass these limitations, super-resolution 3D STED microscopy with 185 nm slice in Z direction was implemented (Fig. 3E) and surprisingly revealed that viral M clusters formed similar “ring-like” structures in close proximity to the “actin rings”. Transmission electron microscopy (TEM) slices of the infected cells also show vesicular structures, of 0.6 to 1 μm size, full of viruses with particle budding events at the cell membranes (Fig.3E, TEM). By STED 3D, the diameter of the “viral rings” ranged between 0.5 and 2 μm with a mean of 0.95 μm and a STD of 0.27 μm (Fig. 3D, bottom). By superposing both F-actin and virus STED images, it appears that M labelled intracellular organelles (“viral rings”) are surrounded by F-actin nanostructures (“actin rings”) (Fig. 3E). Our results allow the hypothesis of a spatial and functional interplay between F-actin nanostructures and M cluster formation during assembly of SARS-CoV-2 particles, suggesting a stabilization of the viral assembly platforms by F-actin or the need of F-actin for the transport of virus loaded vesicles towards the cell plasma membrane.

**Fig 3.**
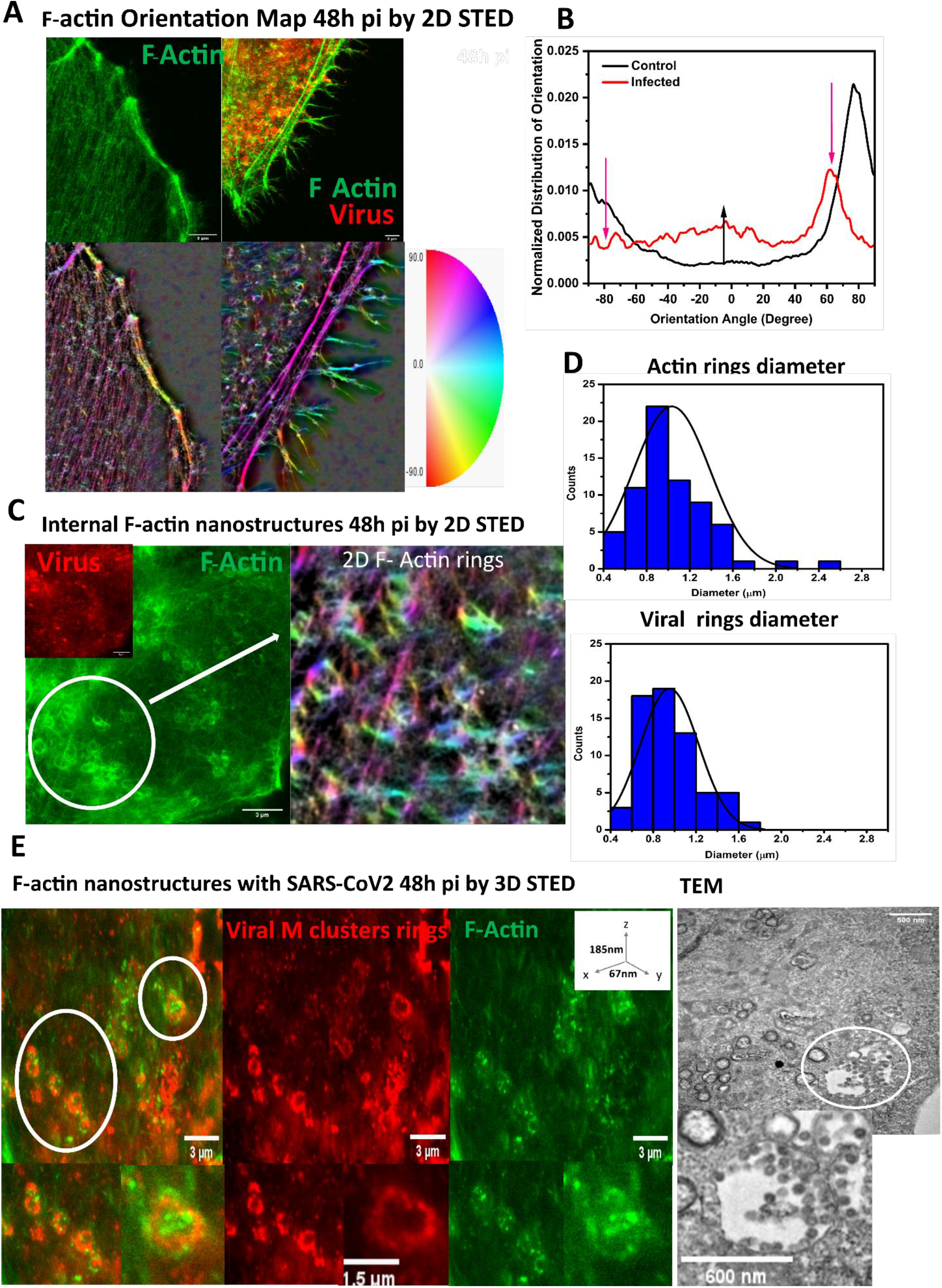
Reorganization of F-actin nanostructures and intracellular actin ring formation in SARS-CoV-2 infected pulmonary cells. STED 2D and 3D images of changes in F-actin and viral clusters of SARS-CoV-2 infected pulmonary cells. A549-hACE2 cells were fixed for control and 48h post infection processed for immunofluorescence coupled to STED microscopy. For imaging virus SARS membrane protein anti-M rabbit antibody and then secondary antibody Star orange (green) and for F-actin Phalloidin Star red (red) were used. (A) 2D STED images and color representation of orientation angle of F-actin network with (48h post infection) or without infection. (B) Plot for distribution of orientation angle, with (48h post infection) or without infection. (C) STED 2D images of actin ring with color representation of rings.(D) Plot for distribution of F-Actin rings and viral ring diameters. (E) STED 3D images of intracellular F-actin rings and viral M clusters in infected pulmonary cells. A number of 25 < *n* <50 cells were analyzed from at least 3 independent experiments. Statistical significant analysis were evaluated using one‐way ANOVA tests. \*\**p* < 0.05. Transmission electron microscopy (TEM) images of intracellular structures filled with budding viruses in SARS-CoV-2 infected pulmonary cells. Scale bars are 500 to 600nm for the zoom image, as indicated.

### SARS-CoV-2 infection induced actin filament rearrangement forming virus loaded protrusions

Production of SARS-CoV-2 particles is a multistep process that occurs in different compartments of an infected cell and culminates in the assembly of virus components at the plasma membrane followed by budding and release of infectious virus particles. To investigate the role of F-actin during virus particle formation and release, we used super-resolution 2D STED microscopy for quantitative analysis (Fig. 4A-C). The number of filopodia-like structures significantly increased from 2 per 10 μm of infected cell plasma membrane (IQR=1-3) in non-infected A549-hACE2 cells to 10 per 10 μm of infected cell plasma membrane (IQR=6-18) in SARS-CoV-2-infected cells 48h pi (Fig. 4B). We also detected a significant increase in the maximum length of filopodia-like structures from 2-4 μM in non-infected cells to 10-12 μm in SARS-CoV-2-infected cells 48h pi (Fig. 4C). Interestingly, we observed these structures being loaded with viruses. A quantitative analysis of the viral M clusters and individuals particles using 2D/3D STED images revealed three different population of viral M clusters in 3 different cellular regions: intracellular (see Fig. 3 D, E), particle release sites at the cell plasma membrane (Fig.4D, zone 1) and viral particles at the filopodia of infected cells (Fig. 4D, zone 2). The size of viral particles ranged in zone 2 from 70 nm to 350 nm with a mean of 209 nm and a STD of 87 nm (Fig. 4D), in zone 1 from 150 nm to 1000 nm (Mean=478 nm; STD=216 nm; Fig. 4D). These quantitative analyses suggested that viral particles are released in package at the cell membrane and then particles migrate (possibly one or two together) on filopodia-like structures. Strikingly, in some infected cells at close proximity, we also observed an inter-connection between cells via filopodia-like structures loaded with viruses, suggesting that virus-containing filopodia could be one of the possible mechanisms for cell-to-cell SARS-CoV-2 infection spread (Fig. 4D).

**Fig 4.**
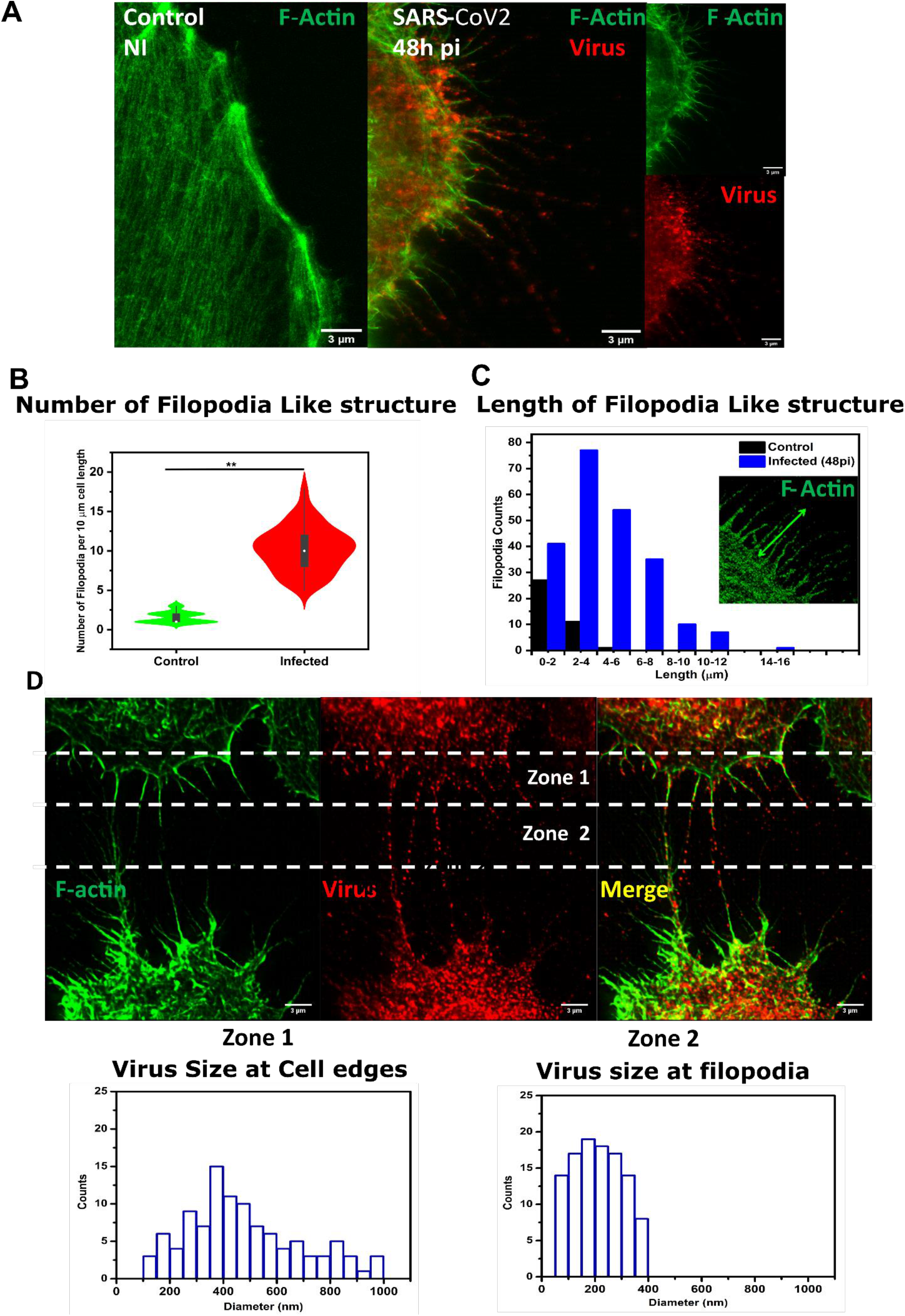
Reorganization of actin fibers into virus loaded filopodia-like protrusions at the cell surface of SARS-CoV-2 infected pulmonary cells. STED 2D images and Quantitative data of changes in F-actin nanostructures in SARS-CoV-2 infected A549-hACE2 cells. A549-hACE2 cells were fixed for control and 48h post infection processed for immunostaining and STED microscopy. For imaging viral particles and clustered SARS membrane protein anti-M rabbit antibody and then secondary antibody Star orange (green) and for F-actin Phalloidin Star red (red) were used. (A) Merge STED 2D Images of control and 48h post infected pulmonary cells. (B) Plot for distribution of number of filopodia per 10 micrometer lengths of each cell infected and control. (C) Plot for distribution of length of individual filopodia in control and infected cells. (D) 3D projection Image and plots showing viral M cluster size at cell edge (Zone 1), at filopodia-like structures (Zone 2), and at cell-to-cell connections. A number of 15 < *n* < 20 cells were analyzed from at least 3 independent experiments. Statistical significant analysis were evaluated using one‐way ANOVA tests. \*\**p* < 0.05.

### SARS-CoV-2 infection enhances expression of actin cytoskeleton regulating genes

Our results suggest that F-actin rearrangements during viral particle assembly are important for viral replication during SARS-CoV-2 infection progression. To gain further insights, we performed a RNA-seq-based transcriptome analysis in non-infected and SARS-CoV-2 infected A549-hACE2 cells at 48h pi (Fig. 5A and Supplementary Fig. 3A). We detected increased levels after infection in 9.91% of the transcripts mapped to the human genome (4376 transcripts with FC ≥ 2), whereas the levels of 7.10% of the transcripts mapped to the human genome were reduced (3136 transcripts with FC ≤ 0.3). Gene set enrichment analysis (GSEA) based on Reactome [23] from the top 9.91% of the genes with increased levels 48h pi (4376transcripts; Fig. 5B) revealed significant enrichment of genes related to RHO GTPases activate PKNs (P= 0.368) as the top item of the ranked list. In addition, graphical representation of the enrichment profile (Fig. 5C) showed a high enrichment score (ES) of 0.559 for RHO GTPases activate PKNs. Since RHO GTPases are best known for their roles in regulating cytoskeletal rearrangements, we monitored the transcript levels of various proteins related to the cytoskeleton (Fig. 5D) and detected increased transcript levels of proteins that are known to be regulated by RHO GTPases, thereby being Alpha-actinin superfamily, and in particular Alpha-Actinins 2 and 3 (ACTN2, ACTN3), the ones with the most prominent increase. Type II myosins are also reported (MHY7 and MHY6), suggesting together with ACTN2 and ACTN3 a possible contractile activity of the infected cells (24), as previously suggested by our cell morphology analysis (Fig. 2). In addition, 2D STED microscopy images and actin orientation analysis (Fig. 3A) revealed the formation of large actin fibers near the cell plasma membrane. Furthermore, WB of protein extracts showed a 2-fold increase of alpha-actinins (ACTN) in SARS-CoV-2 infected cells 48h pi when compared to non-infected cells (Fig. 5E).

**Fig 5.**
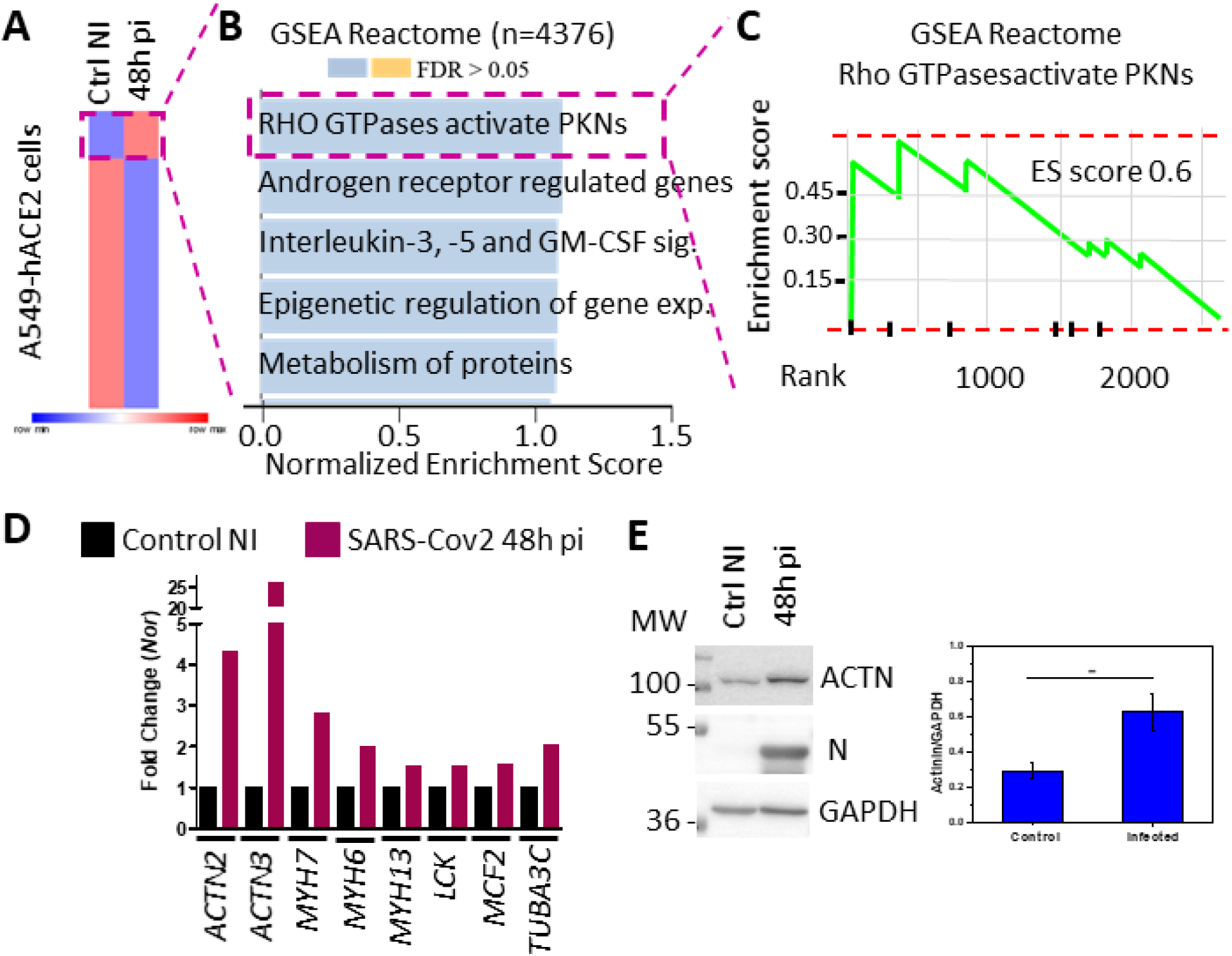
Cellular gene expression analysis of SARS-CoV-2 infected human pulmonary cells using RNAseq reveals an upregulation of alpha-Actinins. (A) Heat map showing RNA-seq-based expression analysis of differentially expressed transcripts in non-infected (Ctrl) and A549hACE2 cells infected with SARS-CoV-2 for 48h (48h pi). *n*=44141 differentially expressed transcripts; 2 individual cell replicates per condition. (B) *Top*. Reactome-based Gene Set Enrichment Analysis (GSEA) of candidates with FC≥2 (*n*=4376 upregulated transcripts) using WebGestalt (WEB-based Gene SeT AnaLysis Toolkit, 2019). *Bottom*. Panther-based Gene Set Enrichment Analysis (GSEA) of candidates with FC≤0.3 (*n*=3136 downregulated transcripts) using WebGestalt (WEB-based Gene SeT AnaLysis Toolkit, 2019). FDR: False Discovery Rate. (C) Reactome-based Gene Set Enrichment Analysis (GSEA) for Rho GTPAses pathway of candidates with FC≥2. (D) Histogram plots representing the basal transcription activity (48h pi normalized to Control) of components of the Rho-GTPase pathway that are differentially expressed in Control and SARS-CoV-2 infected A549hACE2 cells.

### PKN inhibitor reduced SARS-CoV-2 replication in human pulmonary cells

Following the line of ideas from our previous results (Fig. 5, 2 and 3), we investigated the effect of the 2 inhibitors Rho/SFR and PKN inhibitors on SARS-CoV-2 replication, since they regulate actin fibers formation and alpha-actinins regulation, respectively. Data reveals a reduction of SARS-CoV-2 replication in human pulmonary cells overtime and in a dose-dependent manner with an IC50 of 1.36 μM and 0.65 μM for Rho/SFR and PKN inhibitors, respectively (Fig. 6A, Supplementary Fig. 4). The LD50 being determined as 7.02 μM for Rho/SFR inhibitor and 37.7 μM for PKN inhibitor, indicating a selectivity index > 10, thereby supporting that these are potent antiviral inhibitors for the development of therapeutic strategies against SARS-CoV-2. Moreover, confocal immunofluorescence images of human pulmonary A549-hACE2 cells infected with SARS-CoV-2 in the presence of 0.5μM PKN inhibitor showed a restoration of cell morphology and of F-actin structural pattern (Fig. 6B). Strikingly, PKN inhibitor treatment of SARS-CoV-2-infected pulmonary cells reduced the size of intracellular viral M clusters (Fig. 6C) and decreased the levels of viral particle release (Fig. 6D). We used for these experiments remdesivir (IC50 equal 1 μM) as positive control [25], since it is an antiviral drug that targets the virus replication complex reducing the number and size of viral assembly M clusters (Fig. 6C) and decreasing the levels of viral particle release (Fig. 6D). Interestingly, PKN inhibition blocked M clusters in the ER (Supplementary Fig. 5), thus probably at the level of virus assembly or virus egress from the ER, as shown by immunofluorescence staining for M and grp78 ER marker (Supplementary Fig. 5) and suggesting a role for the alpha-actinins superfamily proteins in SARS-CoV-2 assembly and particle egress.

**Fig 6.**
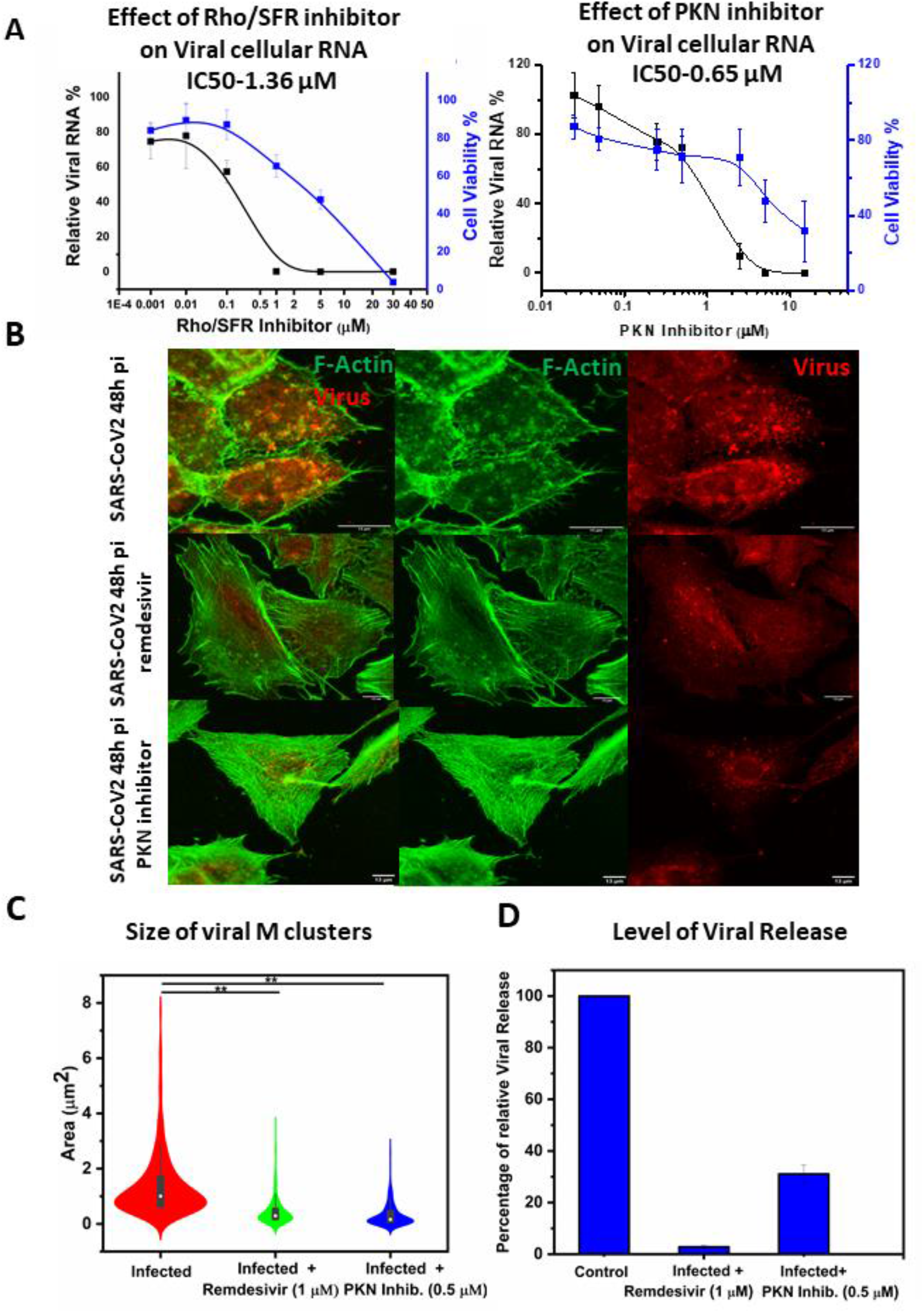
Reduction of SARS-CoV-2 replication in pulmonary cells upon PKN and Rho/SFR inhibitors treatment accompanied with cellular F-actin and cell shape restoration. (A) Dose effect of Rho/SFR and PKN inhibitors on SARS-CoV-2 replication in pulmonary cells using qRT-PCR and cell viability. (B) Confocal images of changes SARS-CoV-2 infected A549-hACE2 cells with the treatment of Remdesivir (1μM) or PKN Inhibitor (0.5 μM). (C) Plot for changes in viral clustered size with or without (infected) treatment of Remdesivir (1μM) or PKN Inhibitor (0.5 μM). (D) Plot for Number of copies of RNA/μL in the supernatant of infected cells at 48h post infection. A549-hACE2 cells were fixed at 48h post infection and processed for immunofluorescence and laser confocal light microscopy using a SARS-CoV-2 membrane protein anti-M rabbit antibody and a secondary antibody Alexa Fluor 568 (in red) and for F-actin imaging Phalloidin Alexa flour 488 (Green) were used. A number of 15 < *n* < 20 cells were analyzed from at least 3 independent experiments. Statistical significant analysis were evaluated using one‐way ANOVA tests. \*\**p* < 0.05.

## Discussion

SARS-CoV-2 is a recently discovered virus. Despite the fact that it is in the spotlight of the scientific community worldwide for being the main cause of the COVID-19 pandemic, the role of the cytoskeleton during SARS-CoV-2 replication has remained elusive. Here, we implemented confocal and super-resolution 2D and 3D STED microscopy to analyse the correlation between that host cell F-actin content and the kinetic of SARS-CoV-2 infection in human pulmonary cells: intracellular SARS-CoV-2 assembly and F-actin content peak at 48h pi, accompanied with a cell morphology deformation and SARS-CoV-2 particle release peaking at 72h pi. The RNA-seq-based analysis of viral transcripts in these infected human pulmonary cells (Supplementary Fig. 1C) correlated with the peak of SARS-CoV-2 assembly at 48h pi. The apparent discrepancies between our SARS-CoV-2 transcriptome and the recently published SARS-CoV-2 transcriptome (26) could be explained by differences in the experimental design, such as the implementation of different cell lines (Calu-3, Caco-2 and Vero cells versus A549-hACE2) and higher viral titters (MOI 1 and 0.1 versus 0.01), among others. The higher susceptibility to SARS-CoV-2 infection of the here implemented human pulmonary A549-hACE2 cells allowed us to use lower viral titers during our RNA-seq experiment (MOI 0.01). More relevant from the present study, the observed kinetics of M cluster formation during SARS-CoV-2 particle assembly and release correlated with rearrangements of cytoskeletal F-actin nanostructures and morphological changes of SARS-CoV-2-infected A549-hACE2 cells. Our results agree with previous reports demonstrating that actin rearrangements are involved during the replication of various viruses targeting the respiratory tract, including the respiratory syncytial virus (RSV) and influenza virus [27,28,29,30]. Further, the strong cytopathic effect observed after SARS-CoV-2 infection could be related with the dramatic changes in F-actin nanostructures and cell morphology. However, the expression of caspases as apoptosis markers were reduced in SARS-CoV-2 infected cells at 48h pi, supporting the viability of the cells analyzed here (Supplementary Fig. 3). It is well known that actin polymerization has a role in replication of influenza viruses (31). For RSV, actin was not completely proven to be involved in assembly, rather actin was playing a role in virus spread driven filopodia induction through Arp2/3 complexes [28]. In our study, increase in F-actin content during infection, with a global actin content remaining quite constant (Figure 1 G), included a redistribution of F-actin polymerization into new nanostructures that appears quite crucial for the late phases of viral replication. Remarkably, we found that SARS-CoV-2 infection induces ring-like F-actin nanostructures surrounding also ring-like viral M-containing structures, suggesting the formation of large intracellular viral organelles, in which SARS-CoV-2 particle assembly take place. This interpretation of our results is in agreement with earlier discoveries, showing that large intracellular structures at the ERGIC containing SARS-CoV-2 structural proteins (M, N, E and S proteins) together with viral genomic RNA and driving the assembly of new viral particles (7,8). Strikingly, we also found that SARS-CoV-2 infection promote filopodia-like structures loaded with viruses to neighbour cells, suggesting these structures as mechanism for cell-to-cell SARS-CoV-2 infection spread. Consistent with these findings, viral cell-to-cell transmission has been reported for other RNA enveloped viruses, such as the human immunodeficiency virus 1 (HIV-1) (32). Summarizing, all these results support a spatial and functional interplay between F-actin nanostructures and M cluster formation during assembly of SARS-CoV-2 particles, which needs to be further investigated.

Actin polymerization starts with the formation of a small aggregate consisting of three actin monomers. Actin filaments are then able to grow by the reversible addition of monomers to both ends. However, one end (the plus end) growth up to ten times faster than the minus end. The polymerisation of actin and the reorganization of actin filaments are complex processes regulated by different factors. One of these factors is the protein kinase N (PKN), which is a fatty acid-activated serine/threonine kinase, whose catalytic domain exhibits homology with that of the protein kinase C family. It has been reported that the interaction of PKN with alpha-actinins is promoted by phosphatidylinositol 4,5-bisphosphate (33), suggesting that PKN/alpha-actinin complexes locate at the cell plasma membrane to promote cortical actin fiber rearrangement (34). Alpha-actinins belong to the spectrin gene superfamily which represents a diverse group of cytoskeletal proteins. Alpha-actinins are F-actin-crosslinking proteins found in various subcellular localizations both in muscle and non-muscle cells (35), and being involved in diverse cellular processes. Besides the involvement of alpha-actinins regulating cortical actin dynamics during HIV-1 entry (36), the role of alpha-actinins during replication of virus has remained largely unknown. Here, we showed that SARS-CoV-2 infection increased levels of *ACTN2* and *ACTN3* transcripts of ACTN proteins. Furthermore, interfering with alpha-actinins function through PKN inhibitor treatment in SARS-CoV-2 infected cells restored F-actin structures and reduced SARS-CoV-2 replication. Overall, our results reveal that F-actin nanostrucutres and F-actin rearrangement are required for SARS-CoV-2 replication in pulmonary host cells and support the idea of using PKN inhibitors for the development of therapeutic approaches against SARS-CoV-2 infection.

## Materials and Methods

### Cell culture and infection

Human pulmonary Alveolar A549-hACE2 cells were obtained from original A549 (ECACC) transduced with a lentiviral vector expressing human ACE2 receptor (manufactured by FlashTherapeutics company, Toulouse, France) and sorted by cytometry for having more than 80% hACE2 on their surface. The sorted A549-hACE2 cells were maintained in RPMI supplemented with 10% heat inactivated fetal bovine serum (FBS), 1% sodium Pyruvate, 0.5% HEPES and antibiotics (penicillin/Streptavidin) and cultivated at 37°C with 5% CO_2_. For virus production, VeroE6 cells were obtained from (ECACC) and maintained in Dulbecco’s minimal essential medium (DMEM) supplemented with 10% heat inactivated fetal bovine serum (FBS) at 37°C with 5% CO_2_.

The strain BetaCoV/France/IDF0372/2020, was supplied by the National Reference Center for Respiratory Viruses hosted by Institut Pasteur (Paris, France) and headed by Pr. Sylvie van der Werf. The human sample from which strain BetaCoV/France/IDF0372/2020 was isolated has been provided by Dr. X. Lescure and Pr. Y. Yazdanpanah from the Bichat Hospital, Paris, France. Moreover, the BetaCoV/France/IDF0372/2020 strain was supplied through the European Virus Archive goes Global (EVAg) platform, a project that has received funding from the European Union’s Horizon 2020 research and innovation program under the grant agreement No 653316. COV-2 Virus was propagated in VeroE6 cells with DMEM containing 2.5% FBS at 37°C with 5% CO_2_ and harvested 72 hours post inoculation. Virus stocks were stored at -80°C and titer using plaque assays as previously described (11).

### Quantitative reverse transcription polymerase chain reaction (qRT-PCR)

RNAs from mock infected or infected (MOI=0.01) A549-hACE2 cell culture supernatant were extracted using the Nucleospin Dx Virus RNA purification kit (Macherey-Nagel). Then qRT-PCR was performed in triplicate as described^20^, using primers targeting the E gene of SARS-CoV-2 (E_Sarbeco-Forward ACAGGTACGTTAATAGTTAATAGCGT; E_Sarbeco-Reverse ATATTGCAGCAGTACGCACACA) and Luna Universal One-Step qRT-PCR Kit (New England Biolabs) on a Roche Light Cycler 480. The calibration of the assay was performed with a nCoV-E-Sarbeco-Control Plasmid (Eurofins Genomics).

### RNA sequencing and data analysis

RNA was sequenced as previously described [PMID 31110176; 29867223]. Briefly, total RNA from non-infected (Ctrl) or SARS-CoV-2 infected A549-hACE2 and hPCLS was isolated using Trizol (Invitrogen). RNA was treated with DNase (DNase-Free DNase Set, Qiagen) and repurified using the miRNeasy micro plus Kit (Qiagen). Total RNA and library integrity were verified on LabChip Gx Touch 24 (Perkin Elmer). One μg of total RNA was used as input for SMARTer Stranded Total RNA Sample Prep Kit-HI Mammalian (Clontech). Sequencing was performed on the NextSeq500 instrument (Illumina) using v2 chemistry with 1x75bp single end setup. Raw reads were visualized by FastQC to determine the quality of the sequencing. Trimming was performed using trimmomatic with the following parameters LEADING:3 TRAILING:3 SLIDINGWINDOW:4:15 HEADCROP:4, MINLEN:4. High quality reads were mapped using with HISAT2 v2.1.0 with reads corresponding to the transcript with default parameters. RNA-seq reads were mapped to human genome hg19. After mapping, Tag libraries were obtained with MakeTaglibrary from HOMER (default setting). Samples were quantified by using analyzeRepeats.pl with the parameters (hg19 -count genes – rpkm; reads per kilobase per millions mapped).

### RNA-seq based expression analysis of viral transcripts

Fastq files from infected A549-hACE2 cells after 24h, and 48h were used as input; for each timepoint we used 2 replicates. Read trimming was performed using trimmomatic (v 0.39) with the following parameters “ILLUMINACLIP:all_adapters_v0.38.fa:2:30:10 AVGQUAL:30 LEADING:0 TRAILING:0 SLIDINGWINDOW:6:30 MINLEN:38”. Trimmed reads where then aligned to the SARS-CoV-2 reference genome NC_045512.2.fasta (downloaded May 2021 from https://www.ncbi.nlm.nih.gov/nuccore/NC_045512) using the STAR (v 2.7.9a) aligner; STAR parameters where the following “--outFilterType BySJout --outFilterMultimapNmax 20 --alignSJoverhangMin 8 --outSJfilterOverhangMin 12 12 12 12 --outSJfilterCountUniqueMin 1 1 1 1 --outSJfilterCountTotalMin 1 1 1 1 -- outSJfilterDistToOtherSJmin 0 0 0 0 --outFilterMismatchNmax 999 -- outFilterMismatchNoverReadLmax 0.04 --scoreGapNoncan -4 --scoreGapATAC -4 -- chimOutType WithinBAM HardClip --chimScoreJunctionNonGTAG 0 –alignIntronMin 20 --alignIntronMax 1000000 --alignMatesGapMax 1000000 -- alignSJstitchMismatchNmax -1 -1 -1 -1”. Samtools (v 1.12) was used to handle the alignments, and bedtools coverage was used to count reads in each viral feature (gene) using the genomic coordinates from GCF_009858895.2_ASM985889v3_genomic.gff (downloaded May 2021 from https://ftp.ncbi.nlm.nih.gov/genomes/all/GCF/009/858/895/GCF_009858895.2_ASM985889v3/GCF_009858895.2_ASM985889v3_genomic.gff.gz) only for protein coding features. Feature counts were transformed to reads per kilobase million (RPKM) we calculated mean RPKM from the duplicates for each feature and then calculated a Fold Change as mean RPKM at 48h / mean RPKM at 24h; this data handling and plotting was performed using R.

### Western Blot analysis

A549-hACE2 cells were infected for 2 hours with SARS-CoV-2 (MOI = 0.01). At different time point (6h, 24h, 48h, 54h, 72h and 77h) post infection (pi), cells were washed twice in PBS, detached with Versen (0.1M EDTA), pelleted at 250g for 6min and lysed in RIPA buffer for western blot analysis. Total protein concentration was calculated using a Bradford protein assay kit (ThermoFisher). 20μg of total cell lysates were diluted in Laemmli buffer and proteins were separated by SDS-PAGE on 8% (for COV-2 S- and N proteins) and 12% (for COV-2 M protein) acrylamide gels. Gels were transferred to PVDF membrane using wet transfer with Tris-glycine-methanol buffer. Membranes were washed in TBS, blocked with 5% milk in TBS-Tween 0.1% for 30min and incubated overnight at 4°C with primary antibodies against the spike S protein (Gentex, cat# GTX632604), N-protein (Gentex, cat# GTX632269) or M-protein (Tebu, cat# 039100-401-A55), all three diluted at 1:1000 in TBS-T. After washing with 5% milk in TBS-Tween, the membranes were incubated with HRP conjugated anti-mouse antibodies for N and S protein, and with HRP conjugated anti-rabbit antibody for M protein and alpha-actinins for 2h at room temperature, then washed in TBS-Tween buffer, incubated with ECL reagent (Amersham cat#RPN2236) and imaged using a Chemidoc Imager (Biorad).

### Immuno-fluorescence confocal and 2D/3D STED super-resolution microscopy

A549-hACE2 cells seeded on glass coverslips were infected with SARS-CoV-2 at a MOI=0.1 or MOI=0.01 (low multiplicity of infection). At different time interval from 6h to 77h post-infection cells were washed with PBS and fixed in 4% paraformaldehyde in PBS for 15 minutes at room temperature, followed by permeabilization with 0.2% Triton X-100 in PBS for 4-5 minutes and blocking in 2% BSA in PBS for 15 min. Incubation with primary antibodies anti-SARS-CoV2 rabbit membrane (M) protein (1:100) was performed for 2 hours at room temperature. After washing with PBS, cells were incubated with secondary antibodies AF568-labeled goat-anti-rabbit (1:200) and Star orange for high resolution STED imaging (1:100) as well as AF488-labeled Phalloidin and Star red phalloidin (1:100) (for high resolution STED microscopy) for 2 hours at room temperature. We have used mounting media prolong gold antifade reagent with DAPI and prolong gold antifade reagent without DAPI for confocal and STED microscopy respectively. Confocal fluorescence images were generated using a LSM800 confocal laser-scanning microscope (Zeiss) equipped with a 63X, 1.4 NA oil objective and STED 2D and 3D measurements were performed on the Abberior Instrument Expert Line STED super-resolution microscope (Abberior Instruments GmbH, Göttingen, Germany) using Star orange 580 and Star red pulsed excitation laser sources with a pulsed STED laser operating at 775 nm. For STED 2D (25% laser) lateral resolution was 67nm and for STED 3D (30% laser) resolution was 185nm in Z. All the images processed with ImageJ/Fiji. For 3D-reconstruction of confocal images, cells were fixed and stained as indicated and imaged as z stack with 0.3 μm sections. Z stack was processed using ImageJ/Fiji, Imaris viewer.

### Electron Microcopy

A549-hACE2 pulmonary cells infected with SARS-CoV-2 were fixed with 2,5% (v/v) glutaraldehyde in PHEM buffer and post fixed in osmium tetroxide 1% / K_4_Fe(CN)_6_ 0,8%, at room temperature for 1h for each treatment. The samples were then dehydrated in successive ethanol bathes (50/70/90/100%) and infiltrated with propylene oxide/ EMbed812 mixes before embedding. 70 nm ultrathin cuts were made on a PTXL ultramicrotome (RMC,France), stained with OTE/lead citrate and observed on a Tecnai G2 F20 (200kV, FEG) TEM at the Electron Microscopy Facility COMET, INM, Platform Montpellier RIO Imaging, Biocampus, Montpellier.

### Statistical analysis

Statistical tests were performed using Origin 2021 software. Statistically significant analysis was evaluated using one‐way ANOVA tests. \*\**p* < 0.05. Cell area and cell volume and height were calculated using 3D viewer plugin from Fiji image J. The orientation angle properties of a given region of interest in an image were computed based on the evaluation of the structure tensor in a local neighborhood using the Java plug-in for ImageJ/Fiji (http://imagej.nih.gov/) ‘OrientationJ’. The Materials and Methods section should provide sufficient information to allow replication of the results. Begin with a section titled Experimental Design describing the objectives and design of the study as well as prespecified components.

## Acknowledgments

We thank CEMIPAI UAR3725 service unit for initial advices on SARS-CoV-2 virus production and titration and Aymeric Neyret for the TEM image. The Microscopy STED was done at the Montpellier Imaging Center for Microscopy (MRI). We are very great full to Marie-Pierre Blanchard for initiation to STED microscopy and to Dr Frederic Eghaian from Abberior for the gift of STED compatible secondary antibodies.

## Author contributions

PM, DB performed cell culture, BSL3 infection, viral stock amplification and titer, viral RNA extraction and qRT-PCR, immunoblots. DB participated to the TEM. JS performed immunofluorescence sample preparation, confocal and STED 2D and 3D Microscopy and quantitative analysis. KR performed RNA extraction and sequencing. KR, IA, SG and GB performed RNA sequencing and analyzed RNA-seq data. DM, JS, GB and KR were involved in manuscript writing. JS, GB and DM conceptualized the study, edited the figures and wrote the manuscript. DM supervised the study. DM and GB raised funding for the study.

## Data and materials availability

All data are available in the main text or the supplementary materials.

## Supplementary Materials

see Supplementary Materials document.

