## Supplemental figures for "Reorganization of F-actin nanostructures is required for the late phases of SARS-CoV-2 replication in pulmonary cells"

[1]Institute of Research in Infectiology of Montpellier (IRIM), University of Montpellier, UMR9004 CNRS, Montpellier, France. [2]Université de Lorraine, CNRS, Laboratoire IMoPA, UMR 7365, F-54000 Nancy, France. [3]Univ Paris Est Creteil, Glycobiology, Cell Growth and Tissue Repair Research Unit (Gly-CRRET), Brain and Lung Epigenetics (BLUE), Creteil, France. [4]International Laboratory EPIGEN, Universidad de la Salud del Estado de Puebla, 72308 Puebla, Mexico. [5]Instituto Nacional de Medicina Genómica (INMEGEN), Mexico city, Mexico. [6]ECCPS Bioinformatics and Deep Sequencing, Max-Planck-Institute for Heart and Lung Research, 61231 Bad Nauheim, Germany.

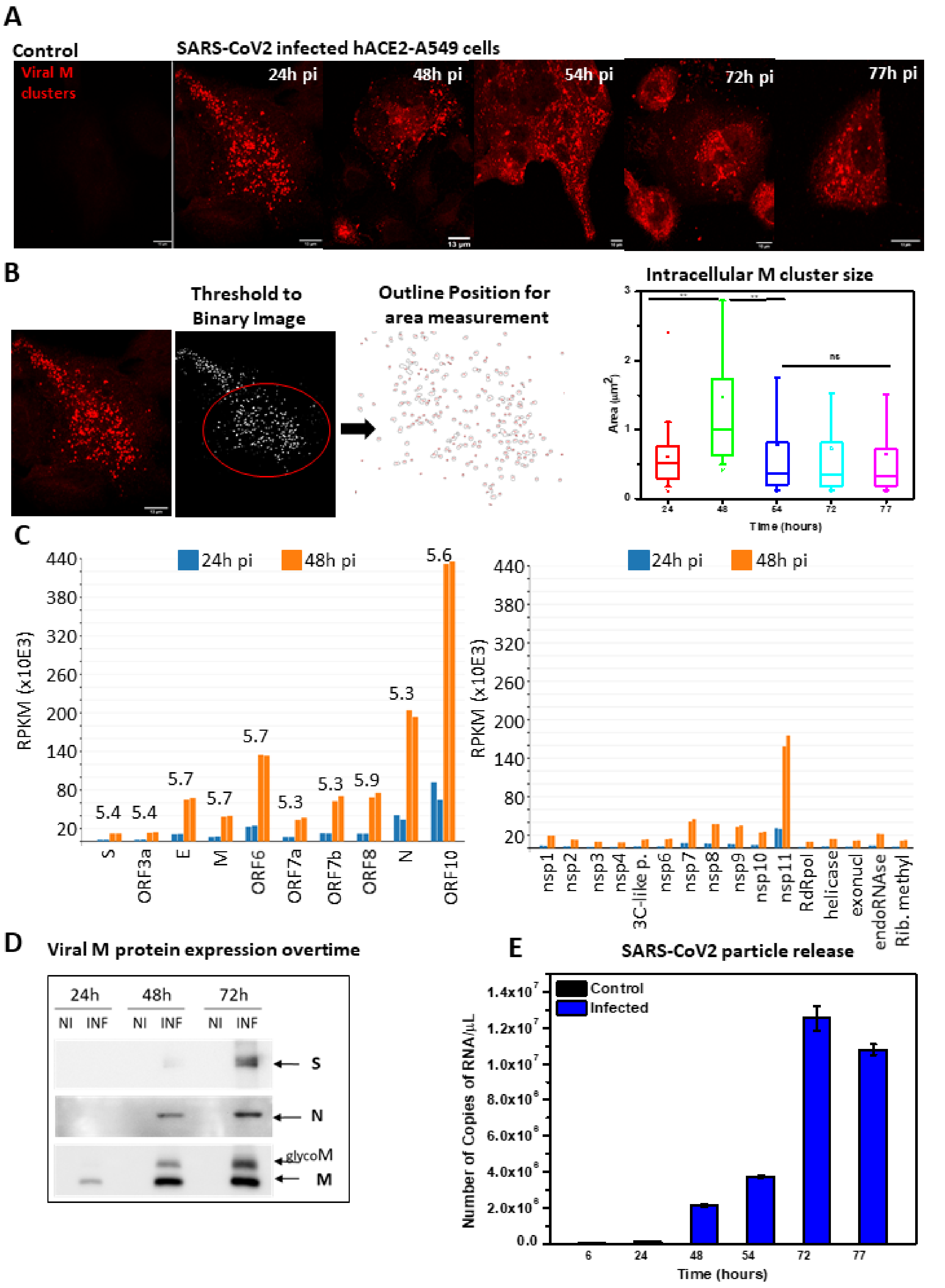


**Figure S1**: **SARS-Cov2 viral assembly clusters growth kinetics, virus release and global actin content upon SARS-CoV2 infection in human pulmonary A549-hACE2 cells**.

Imaging and quantitative analysis of time course changes in viral M clusters area, of SARS-CoV-2 infected A549-hACE2 cells. A549-hACE2 cells were fixed at 0h, 24h, 48h, 54h, 72h and 77h post infection and processed for immunofluorescence. SARS-CoV-2 membrane protein anti-M rabbit antibody and a secondary antibody Alexa Fluor 568 (in red) and for F-actin Phalloidin Alexa Fluor 488 were used for confocal microscopy. (A) Confocal images of viral clusters with different time post infection 0h to 77h. (B) Plot for changes in viral clustered size at different time post infection. A number of $20 < n < 50$ cells were analyzed from at least 3 independent experiments. Statistical significant analysis were evaluated using one-way ANOVA tests. $^{**}p < 0.05$. (C) Plot for RNAseq vial gene expression analysis at different time post infection. D) Western blot data for viral structural protein expression with non-infected and infected cell at different time point post infection. (E) Plot for Number of copies of RNA/μL in the supernatant of infected cells with different time post infection.

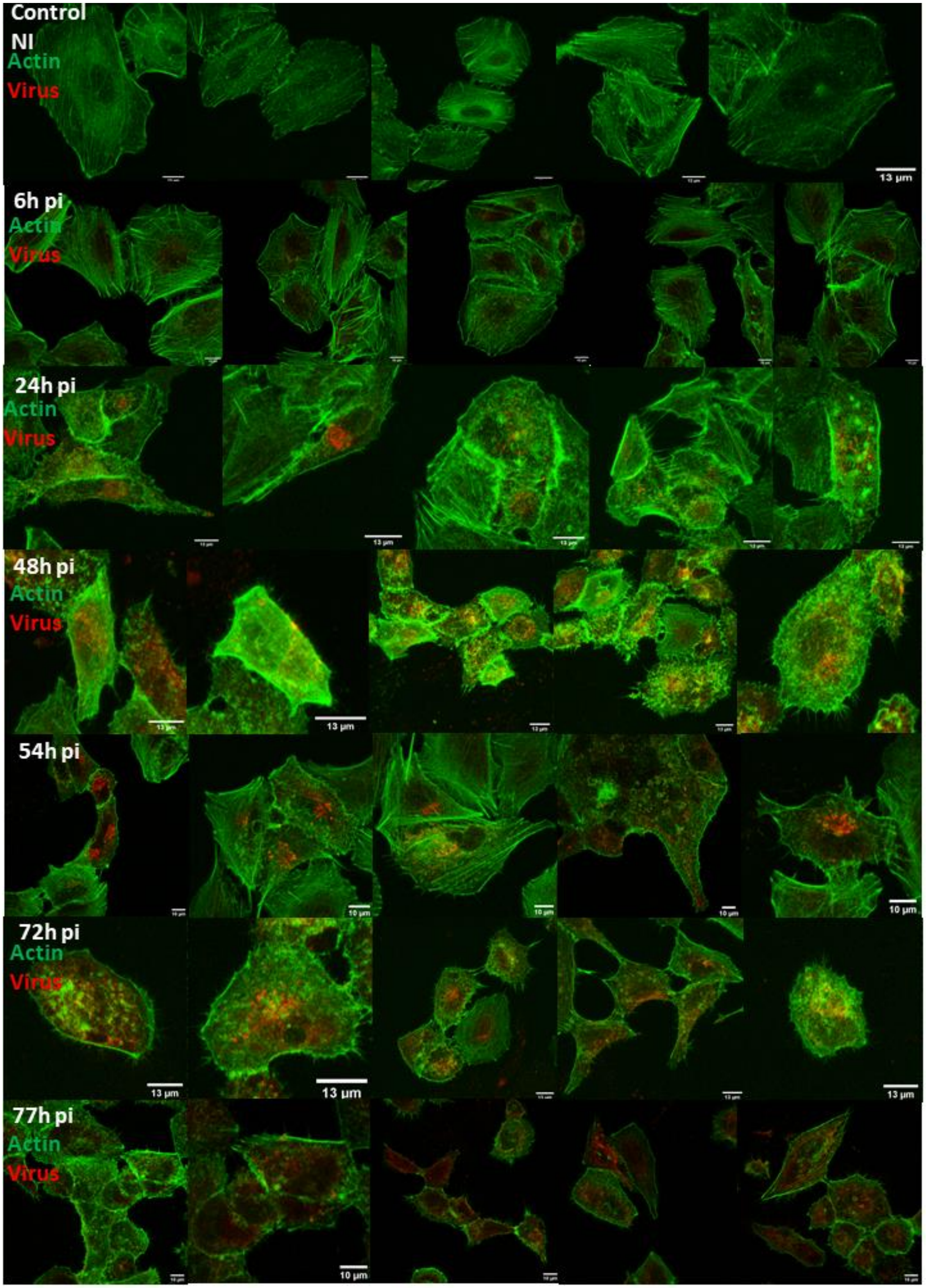


**Figure S2**: **Confocal images of F-actin cytoskeleton and viral clusters overtime upon SARS-CoV2 in human pulmonary cells.**

Library of Images at time course changes in F-Actin and viral M clusters of SARS-CoV-2 infected A549-hACE2 cells. A549-hACE2 cells were fixed at 0h, 24h, 48h,54h, 72h and 77h post infection and processed for immunofluorescence. SARS-CoV-2 membrane protein anti-M rabbit antibody and a secondary antibody Alexa Fluor 568 (in red) and for F-actin Phalloidin Alexa Fluor 488 were used for confocal microscopy.

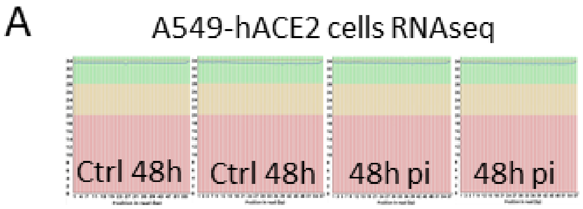


| Sample | Total reads | Surviving reads | Alignment rate (hg19) |
|---|---|---|---|
| Ni 24h a | 25335420 | 25193888 (99.44%) | 92.55% |
| Ni 24h b | 33680299 | 33507122 (99.49%) | 92.73% |
| Inf 24h a | 30959519 | 30795453 (99.47%) | 82.15% |
| Inf 24h b | 31656711 | 31483669 (99.45%) | 83.34% |
| Ni 48h a | 35770502 | 35594957 (99.51%) | 93.01% |
| Ni 48h b | 24233110 | 24113590 (99.51%) | 93.11% |
| Inf 48h a | 42029545 | 41821432 (99.50%) | 36.43% |
| Inf 48h b | 34432891 | 34270524 (99.53%) | 38.17% |

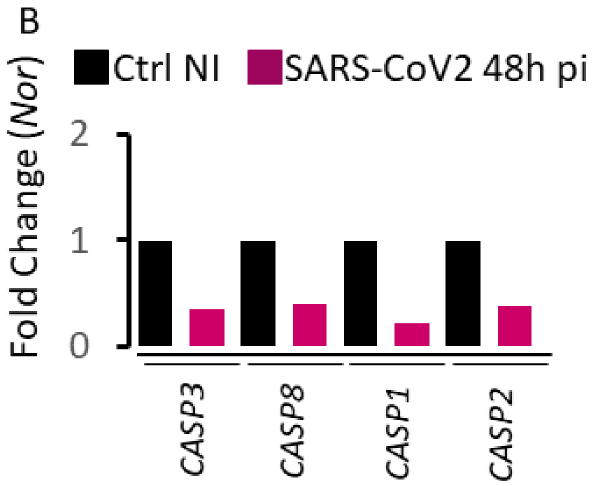


**Figure S3**: **Cellular gene expression of human pulmonary cells SARS-CoV-2 infected and non-infected using RNAseq.** (A) Reproductibility of RNAseq analysis of A549-hACE2 infected cells at 24h and 48h pi and non-infected cells (Ni). (B) Cellular Caspase gene expression at 48h post-infection in human A549hACE2 pulmonary cells using RNAseq: Plot profile of Caspase Fold Change upon infection, non-infected control (Ctrl NI) and SARS-CoV-2 48h pi.

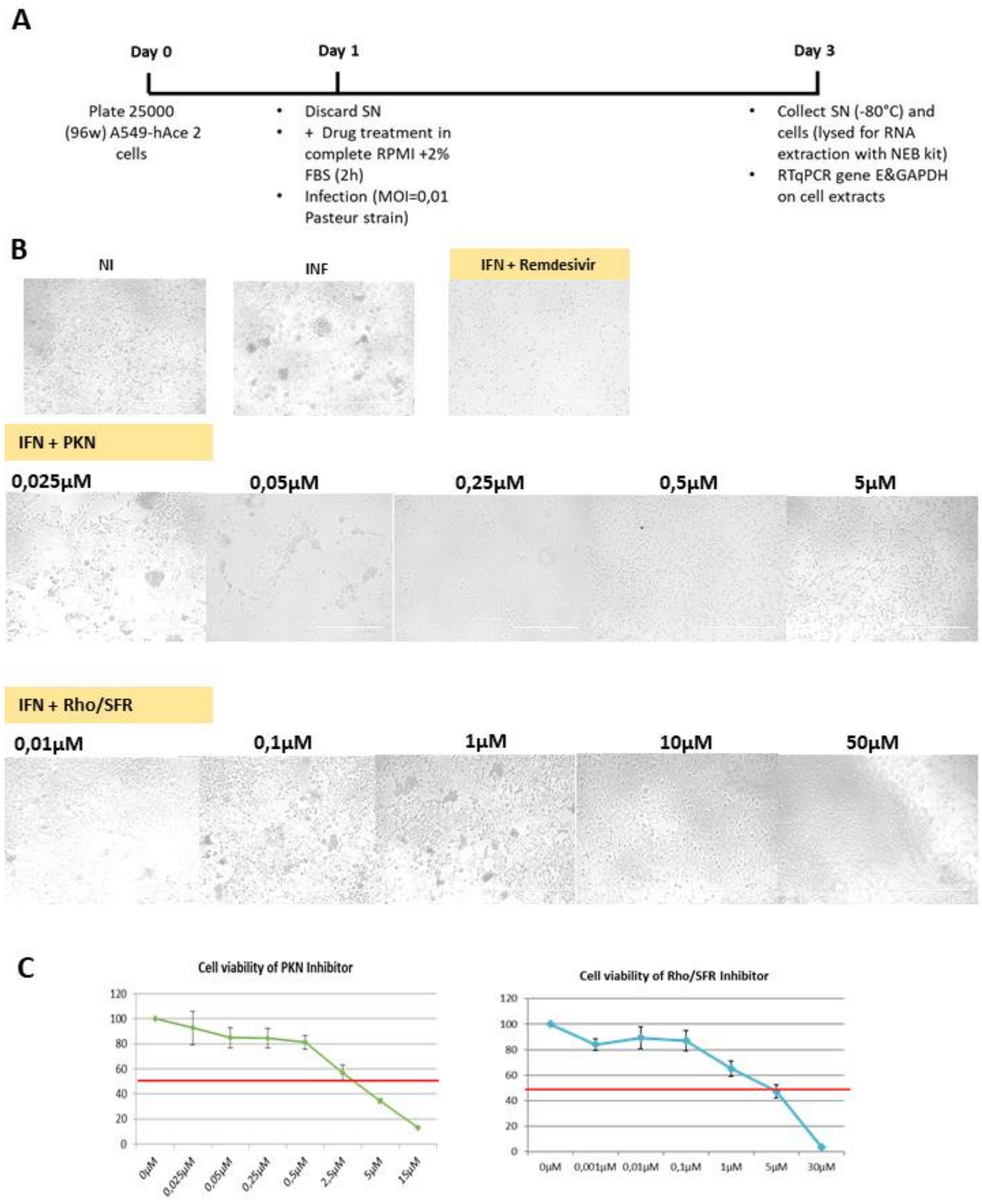


**Figure S4**: **Cytopathic effects of SARS-CoV-2 on human pulmonary A549hACE2 cells**. A) Scheme of drug treatment and infection of A549hACE2 cells. B) Light microscopy images (10x) showing the cytopathic effects of SARS-CoV2 infection on human pulmonary cells in the presence and in the absence of different drugs targeting F-actin signalization under Rho/SFR or PKN/alpha-Actinins regulation. (C) Cell viability in function of drug concentrations.

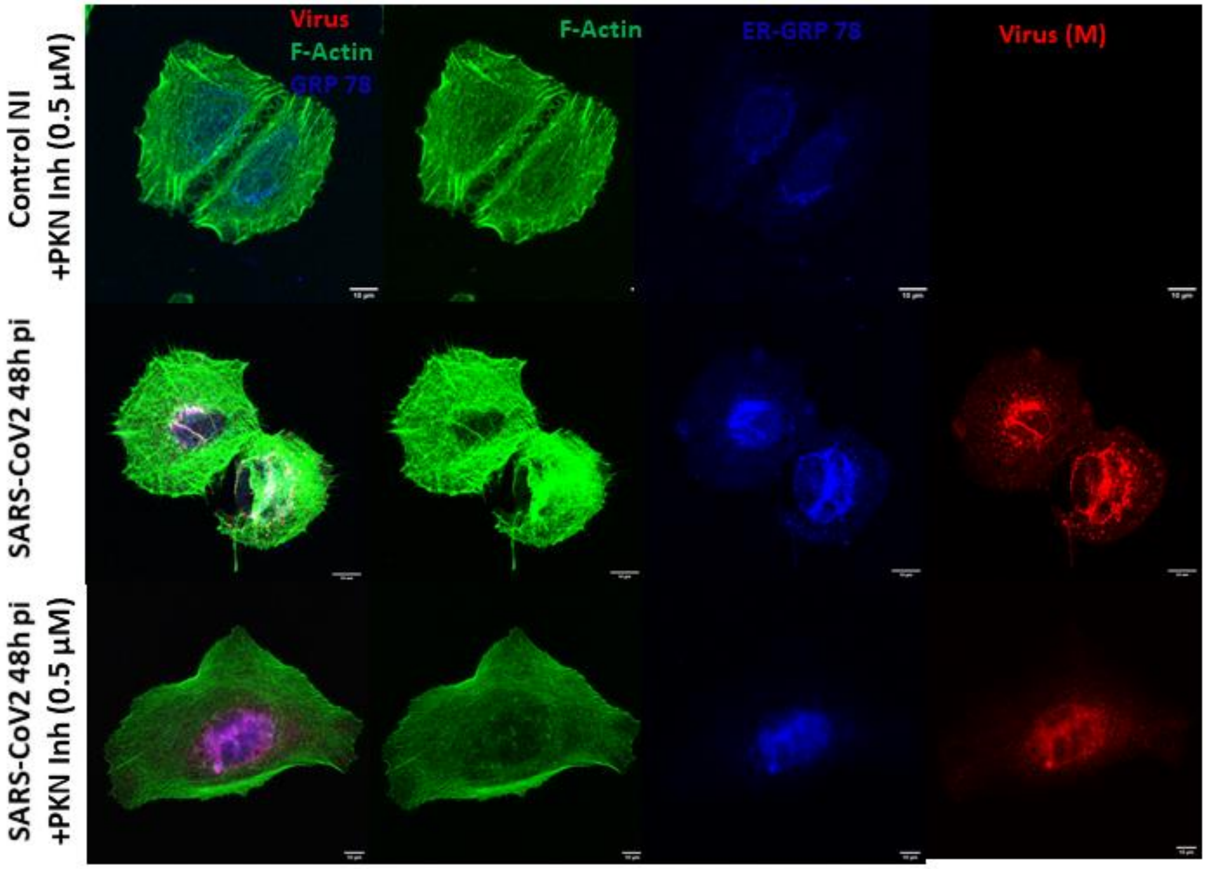


**Figure S5**: Confocal images of F-actin, virus and the Endoplasmic Reticulum (ER) upon SARS-CoV2 infection (48h pi) in human pulmonary cells in the presence and in the absence 0,5µM of PKN inhibitor showing a retention of M clusters in the ER upon drug treatment. F-actin is labelled with Alexa488-Phalloidin (green), ER is labelled with GRP-78 antibody labelled with Alexa G- 633 (blue) and SARS-CoV2 is labelled with anti-M antibody with Alexa R-568 (red). Scale bar is 10µm.

**Other Supplementary Materials for this manuscript include the following:**

**Movie S1: 3D images of non-infected control pulmonary cells**. Human pulmonary A549-hACE2 cells were fixed at 48h and processed for immuno-fluorescence confocal microscopy. For F-Actin imaging Phalloidin Alexa-488 (Green) was used to stain the cells.

**Movie S2: 3D images of SARS-CoV-2 infected pulmonary cells**. Human pulmonary A549-hACE2 cells were fixed at 48h post-infection with SARS-CoV-2 and processed for immuno-fluorescence confocal microscopy. For imaging viral clusters with CoV-2 membrane protein anti-M rabbit antibody and then secondary antibody Alexa Fluor 568 (Red) and F-Actin imaging
